# A midbody component homolog, *too much information/prc1-like*, is required for microtubule reorganization during both cytokinesis and axis induction in the early zebrafish embryo

**DOI:** 10.1101/2021.06.10.447958

**Authors:** S Nair, E.L. Welch, C.E. Moravec, R.L. Trevena, F. Pelegri

## Abstract

We show that the zebrafish maternal-effect mutation *too much information* (*tmi*) corresponds to zebrafish *prc1-like* (*prc1l*), which encodes a member of the MAP65/Ase1/PRC1family of microtubule-associated proteins. Embryos from *tmi/prc1l* homozygous mutant mothers display cytokinesis defects in meiotic and mitotic divisions in the early embryo, indicating that *tmi/prc1l* has a role in midbody formation during cell division at the egg-to-embryo transition. Unexpectedly, maternal *tmi/prc1l* function is also essential for the reorganization of vegetal pole microtubules required for embryonic axis induction. While Prc1 is widely regarded to crosslink microtubules in an antiparallel conformation, our studies provide evidence for an additional function of Prc1 in the bundling of parallel microtubules in the vegetal cortex of the early embryo during cortical rotation and prior to mitotic cycling. These findings highlight common yet distinct aspects of microtubule reorganization that occur during the egg-to-embryo transition, driven by maternal product for the midbody component Prc1l and required for embryonic cell division and pattern formation.

## Introduction

Development prior to zygotic genome activation in animal embryos, including the model system *Danio rerio*, relies on inherited maternal factors (Pelegri 2003, Abrams and Mullins 2009), many of which are RNAs that become translated products after egg activation (Sheets et al. 2017). The process of translation allows to generate a protein supply sufficient for the fast-growing embryo, but also generates a temporal lag in product availability. Because of this lag, the earliest events in embryonic development must be driven by ready-to-function maternal factors, which include already made proteins present in the egg. Since eggs are arrested at meiosis and are transcriptionally quiescent (Svoboda et al. 2017), such maternal products must be synthesized prior to the completion of oocyte maturation. These same maternal products, present in the egg for the early embryo, are also available for use at earlier stages through oocyte maturation and meiotic resumption during egg activation. The temporal usage of such early maternal factors, indicates a process in which the egg-to-embryo transition is not a sharp boundary occurring at egg activation, but instead spans the completion of meiosis, egg activation and fertilization, and the earliest embryonic mitotic cycles (Courtois et al. 2012).

A key event that occurs during meiosis after egg activation and in the early embryo is cell division, a process that relies on extensive cytoskeletal remodeling. Early embryonic cells are unusually large and additionally involve patterning determinants (Lindeman and Pelegri 2010, Wühr et al. 2010), conditions that require unique processes within these cells including dynamic cytoskeletal reorganization. In the early embryo, cytoskeletal changes are particularly evident not only in the spindle apparatus during mitosis but also during furrow formation and maturation (Jesuthasan 1998, Wühr et al. 2010, Eno and Pelegri 2018). At furrow induction, ends of anaphase astral microtubules deliver signals to the cleavage plane to induce furrow formation (Rappaport 1996, Yabe et al. 2009, Nair et al. 2013). These microtubules transition into an array of tubules parallel to each other and perpendicular to the furrow along the length of the furrow, the furrow microtubule array (FMA) (Danilchik et al. 1998, Jesuthasan 1998). During furrow maturation, FMA tubules continue to undergo increasing bundling as they appear to migrate and form a compact structure at each of the furrow distal ends (Pelegri et al. 1999, Eno et al. 2018). The FMA during furrow maturation in the zebrafish has been proposed to be analogous to midbody formation in smaller cell types (Eno et al. 2018), with the FMA exhibiting features similar to those of midbodies, such as microtubule bundling, coupling to the exocytosis of internal membrane vesicles and eventual microtubule disassembly (Jesuthasan 1998, Pelegri et al. 1999, Danilchik et al. 2003, Eno et al. 2018). Factors known to participate in the formation or regulation of the mammalian midbody, such as components of the Chromosomal Passenger Complex and Centralspindlin, are also associated with the FMA (Chen et al. 2002, Yabe et al. 2009, Nair et al. 2013).

As the zygote initiates meiotic resumption and the embryonic cell cycles, other processes essential for embryonic development also ensue. In fish and amphibians, a key event is the off-center displacement of dorsal determinants from the vegetal pole, where they are located during oogenesis, towards their site of action at the future dorsal side of the embryo (reviewed in (Blum et al. 2014, Welch and Pelegri 2017)). As in the case of cytokinesis, this symmetry-breaking event also depends on the reorganization of microtubules, this time at the vegetal cortex (Elinson and Rowning 1988, Houliston and Elinson 1991, Jesuthasan and Strähle 1997, Marrari et al. 2003, Marrari et al. 2004, Tran et al. 2012, Ge et al. 2014). Such vegetal cortex microtubules are thought to undergo rapid polymerization after egg activation, with the resulting microtubules bundling as parallel arrays that run along cortical arcs towards the presumptive dorsal site and which mediate the directed movement of dorsal determinants (Houliston and Elinson 1991, Schroeder and Gard 1992, Marrari et al. 2003, Tran et al. 2012, Olson et al. 2015).

Here we describe a zebrafish maternal-effect mutation *too much information* (*tmi*),originally identified as essential for cytokinesis in the early embryo. We determine that *tmi* corresponds to Protein regulator of cytokinesis 1-like *prc1-like* (prc1l), a protein that is part of the MAP65/Ase1/PRC1 family of microtubule-associated proteins known to control the formation of the midzone and the completion of cytokinesis. We find that zebrafish Tmi/Prc1l functions in cytokinesis during meiotic and early embryonic mitoses, specifically in microtubule network changes involved in spindle formation and FMA reorganization during cell division. We additionally uncover an unexpected role for *tmi* in the reorganization of the microtubule cytoskeleton at the vegetal pole, required for the asymmetric segregation of dorsal determinants in the early embryo. Unexpectedly and contrary to the widely known function of Prc1 as a crosslinker dedicated to antiparallel microtubules at the spindle midzone and midbody, our studies indicate a functional role for Prc1 in the bundling of parallel vegetal cortex microtubules during cortical rotation. Our results show that *tmi/prc1* functions in microtubule bundling during both FMA and vegetal cortex microtubule reorganization, revealing common factors driving contemporaneous processes essential for cell division and pattern formation.

## Results

### A maternal-effect mutation in *tmi* affects cytokinesis in the early embryo

The *tmi^p4anua^* mutation was isolated in an ENU-induced mutagenesis screen for recessive zebrafish maternal-effect genes (Dosch et al. 2004) and results in early cytokinesis defects in the embryo (Nair et al. 2013). Females homozygous for *tmi^p4anu^* develop into viable, phenotypically wild-type adults; however, 100% of embryos from such females (henceforth referred to as *tmi* mutant embryos) display cell division and cytokinesis defects that result in the formation of an acellular embryo (Fig. 1) followed by lysis around 6 hours post fertilization (hpf; data not shown). Processes characteristic of egg activation, such as chorion expansion and ooplasmic streaming leading to the lifting of the blastodisc, which occur during the first 30 minutes post fertilization (mpf), appear normal in *tmi* mutant embryos (Fig. 1A,E, and data not shown). Differences between wild-type and mutant embryos begin to appear upon initiation of cell division. At approximately 45 mpf, wild-type embryos have completed the first cell cycle with a clearly evident furrow leading to a two-cell embryo (Fig. 1A), and cleavage furrows corresponding to subsequent cell cycles occur every 15 minutes thereafter (Fig. 1B-D). In contrast, *tmi* mutant embryos at similar stages exhibit furrows that either fail to form or do not undergo furrow deepening (Fig. 1E-H). Labeling to detect DNA and α-tubulin during early stages of *tmi* mutants (Fig. 1I,J) shows an apparently normal mitosis progression, as assessed by the number of nuclei, and confirms furrow initiation, as suggested by the appearance of a microtubule exclusion zone at furrow sites. However, *tmi* mutant furrows do not accumulate membrane markers such as ß-catenin, a component of the cell adhesion junction present in mature furrows of the early zebrafish embryo (Fig. 1K,L), indicative of defects in furrow completion.

**Figure 1.**
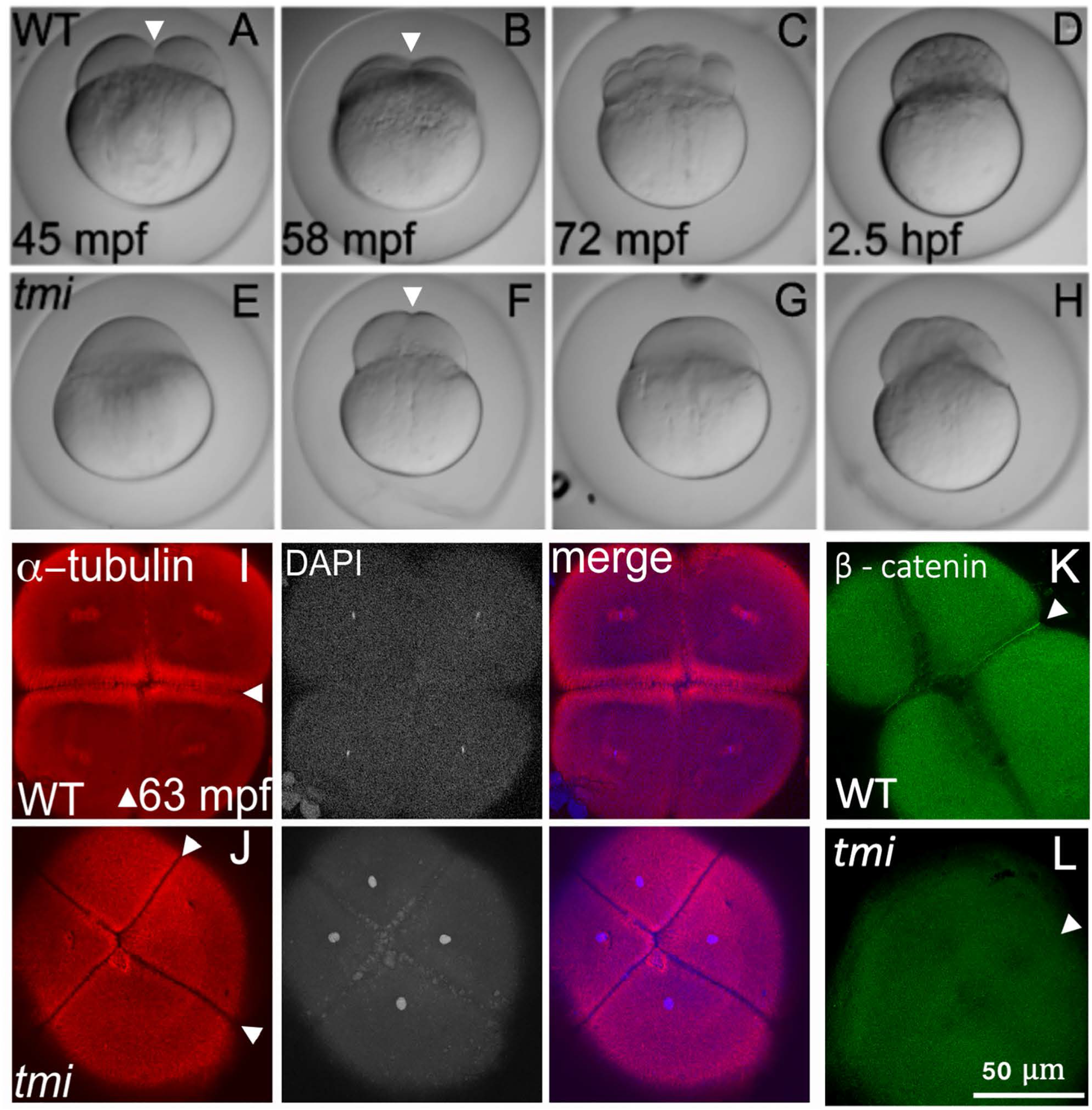
*tmi* mutant embryos exhibit defects in cytokinesis. (A-H) Developmental time course of wild type (A-D) and *tmi* mutant (E-H) embryos. In wild type, furrow formation (arrowheads in (A,B) and cell division proceeds to generate a blastula. In *tmi* mutants, incipient furrows form (arrowhead in F) but do not fully contract during cell division, resulting in acellular embryos. (I-L) Immunolabeling of fixed embryos. (I) In wild type, labeling of microtubules (a-tub) and DNA (DAPI) show formation of furrows (arrowheads) including reorganization of the microtubular apparatus characteristic of maturing furrows (the FMA, arrowheads in I) as well as nuclear division. (J) In *tmi* mutants, furrows appear to initiate development as assessed by zones of microtubule exclusion (arrowheads in J), although the FMA does not fully form, exhibiting a reduction in microtubule enrichment. (K) Wild-type embryos show accumulation of ß-catenin in mature furrows (arrowheads in K), but *tmi* embryos lack ß-catenin accumulation (arrowheads in L). All phenotypes are 100% penetrant with 7-15 embryos per condition in (I-L) (see Methods).

At later stages in mutant embryos, nuclei-like structures appear to accumulate in the absence of membrane formation, with these structures showing aberrant positioning and apparent clumping into unevenly-sized large clusters (Nair et al. 2013, data not shown, see also Supp. Fig 2). These large DNA aggregates are similar to those observed in embryos mutant for other genes and which lack membranes (Pelegri et al. 1999, Nair et al, 2013, Eno et al 2016), and are likely formed through the aberrant attraction of chromatin by neighboring asters in the absence of cellular membranes (Tanaka et al. 2005, Shrestha and Draviam 2013).

### *tmi* encodes Prc1-like, a maternally-expressed protein of the Microtubule-associated protein (MAP65/ASE1) family

Segregation analysis of the *tmi^p4anua^* mutant allele using Simple Sequence Length Polymorphisms (SSLPs) mapped the mutation to a ~20Mb cM region in linkage group 21 (see Methods). Amongst candidate genes in the region, zgc:86764 in the chromosome harboring the mutation contained a potential causative mutation resulting in the conversion of a conserved Leucine at position 221 into a stop codon (Fig. 2A,B; Sup. Fig. 1A). zgc:86764 corresponds to *prc1-like* (*protein regulator of cytokinesis 1-like, prc1l*), belonging to a family of microtubule associated proteins that include MAP65 in Arabidopsis, Ase1 in yeast, and PRC1 in mammals, characterized by the presence of the MAP65/Ase1/PRC1 homology domain involved in microtubule binding (Jiang et al. 1998, Schuyler et al. 2003, Smertenko et al. 2006) and required for spindle midzone assembly during cytokinesis (Mollinari et al. 2002, Eggert et al. 2006, Zhu et al. 2006). Wild-type Prc1l encodes a 580 amino acid protein and the mutated *tmi* allele codes for a severely truncated protein that lacks the C-terminal 350 amino acids, including the majority of the MAP65/Ase1/PRC1 domain, and is therefore likely a functional null.

**Figure 2.**
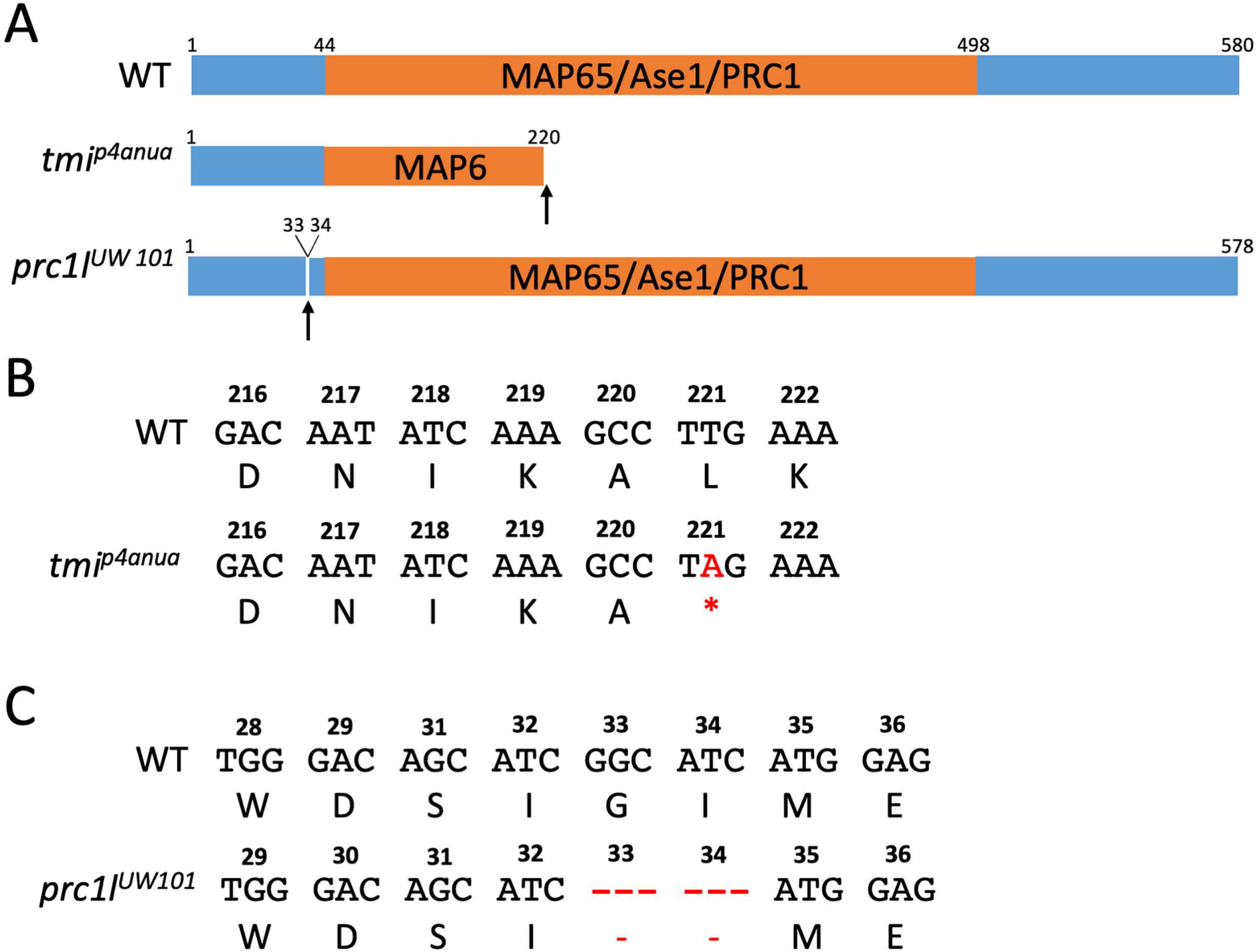
Wild-type and mutant zebrafish PRC1L. (A) Diagram of the predicted wild-type protein, depicting the conserved MAP65/Ase1/PRC1 domain. The lesion in the ENU-induced maternal-effect allele *tmi^p4anua^* results in a truncation of the protein deleting a majority of this domain. The CRISPR-Cas9-induced allele *prc1l^UW101^* results in the deletion of 2 conserved amino acids (see Sup. Fig. 1B and Sup. Fig. 7) N-terminal to the conserved MAP65/Ase1/Prc1 domain. (B-C) Nucleotide sequence and translated amino acids showing the creation of a premature stop codon in *tmi^p4anua^* (B) and the 2-amino acid deletion in *prc1l^UW101^* (C).

To validate the gene assignment of the originally isolated maternal-effect mutation *tmi* as *prc1l*, we utilized CRISPR/Cas-9 to induce independent mutations at the *prc1l* locus, using a guide RNA targeted to a conserved region of the protein upstream of the MAP65/Ase1/PRC1 domain (see Methods). We isolated a six base pair deletion allele *prc1^uw101^* that removed two conserved amino acids (Fig. 2A,C, Sup. Fig. 1B). Female fish transheterozygous for the *tmi^p4anua^* and *prc1^uw101^* alleles exhibited an acellular maternal-effect phenotype indistinguishable from that in embryos from *tmi^p4anua^* homozygous mothers (Sup. Fig. 2), indicating non-complementation between the two alleles and corroborating that *tmi* corresponds to *prc1l*.

*prc1l* is one of three PRC1 gene paralogs in the zebrafish: *prc1a, prc1b* and *prc1l*, with *prc1a* on linkage group 25 and *prc1b* on linkage group 7 (Sup. Fig. 3, Sup. Fig. 4, and data not shown). Analysis of RNA-seq mRNA baseline database available via the Expression Atlas (an ELIXIR database service (http://www.ebi.ac.uk/gxa/experiments/E-ERAD-475)) that provides a baseline transcriptional profiling for multiple developmental stages of the zebrafish, from zygote to 5-day post-fertilization (White et al. 2017), shows high *prc1l* RNA levels early in embryonic development that become significantly reduced at the end of the blastula stage period (dome stage, 4.33 hpf) (Sup. Fig 3A), consistent with *prc1l* functioning as a maternal-specific gene. Whole mount in situ hybridization shows prc1l RNA is evenly distributed in 1-to 4-cell wild-type embryos, with similar distribution in *tmi* mutants (Sup. Fig. 3B). In contrast to *prc1l, prc1a* exhibits expression during early development through the gastrula period up to the early segmentation period (10.33 hpf), while *prc1b* shows low levels of expression throughout early development (Sup. Fig 3A). Phylogenetic analysis comparing *prc* family genes in vertebrate species revealed that zebrafish *prc1a* and *prc1b* are more closely related to a canonical PRC1 present throughout vertebrate species, whereas *tmi/prc1l* clusters closer to a gene duplicate present only in fish and amphibian lineages (Sup. Fig. 4). These data further support the notion that *prc1l* is an ancient gene duplicate that specifically acts at the egg-to-embryo transition.

### *tmi/prc1l* function is required for microtubule dynamics in embryonic cell divisions

Given the known role for Prc1 on midzone formation, we visualized spindles in early embryos using antibodies against α-tubulin and γ-tubulin to label microtubules and centrosomes, respectively, and DAPI to label DNA. In early wild-type embryos during metaphase for the first cell cycle (~45mpf), chromosomes are aligned at the metaphase plate and metaphase astral microtubules emanate from the centrosomes at each spindle pole (Fig. 3A; 14/14; Jesuthasan 1998, Wühr et al. 2010, Wühr et al. 2011)). At this time, mutant embryos show defects in spindle formation, exhibiting instead shorter and wider spindles that are often asymmetric, with a pole larger than the other (Fig. 3B, two detectable spindle poles as ascertained by γ-tubulin localization, 7/20; Fig. 3C, one detectable spindle pole, 13/20). Later at around 60 mpf, chromosomes in wild-type embryos begin movement towards the poles characteristic of anaphase (Fig. 3D, 15/15; Jesuthasan and Strähle 1997, Wühr et al. 2010, Wühr et al. 2011), whereas in *tmi* mutants spindle morphology remains abnormal, continuing to exhibit spindle pole asymmetries and additionally showing mis-segregated chromosomes (Fig. 3E, 23/23 with abnormal spindle morphology, 17/23 with mis-segregated chromosomes). Thus, *prc1l* is essential for proper spindle structure and dynamics during the early blastomere divisions.

**Figure 3.**
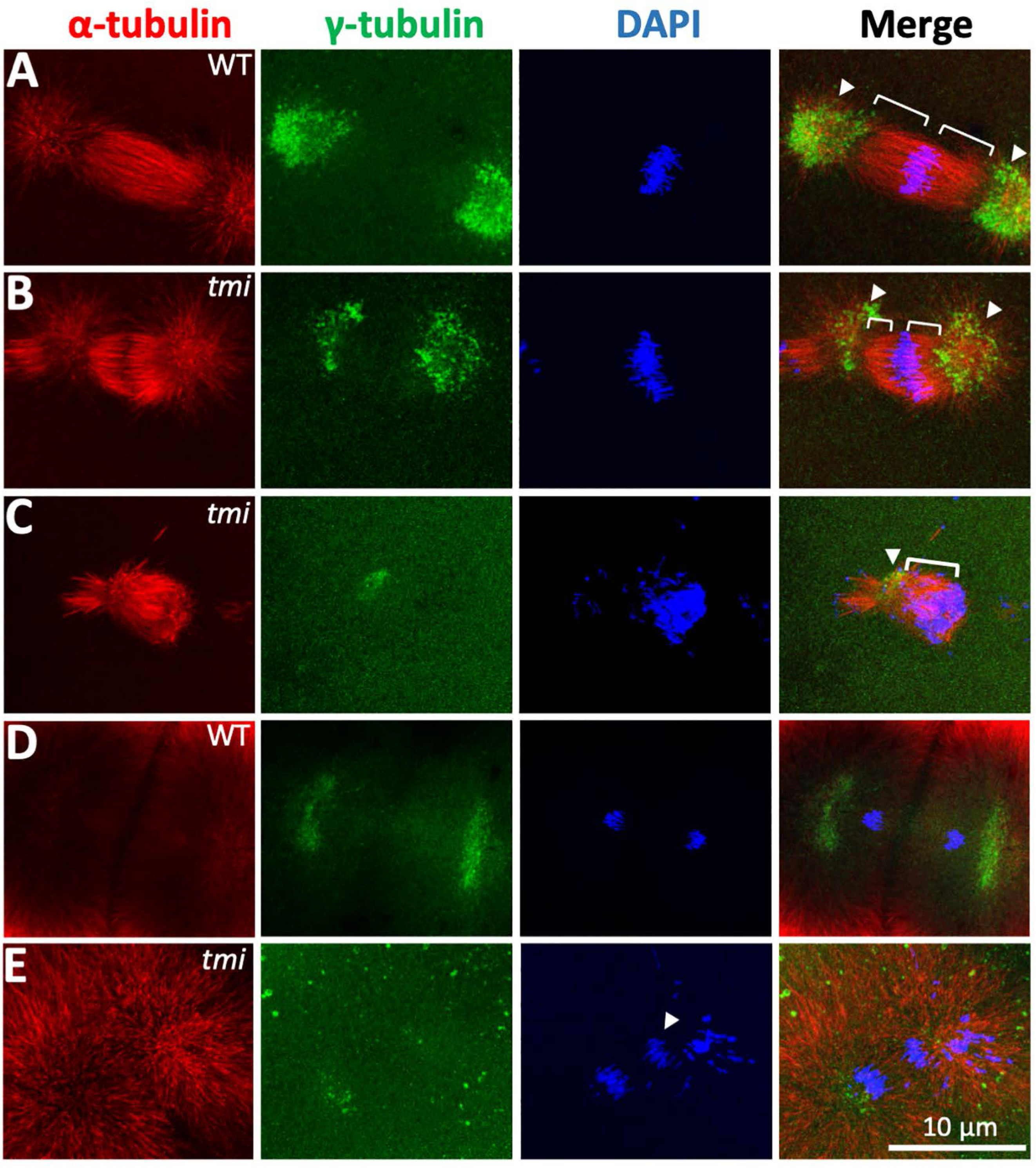
Spindle formation defects in the early mitotic divisions in *tmi/prc1l* mutants. (A-C) Wild-type spindles at metaphase (A), with metaphase astral microtubules colocalizing with the centrosomal marker γ-tubulin. Wild-type asters are highly symmetric, with equal-sized length of kinetochore microtubule and centrosomes. In *tmi/prc1l* mutants, spindles at this same time point are often asymmetric (B) or unipolar with chromosomes clustering at one end of the structure (C). Brackets and arrowheads in (A-C) highlight kinetochore microtubule and centrosomes, respectively. (D-E) During anaphase, *tmi/prc1l* mutants exhibit chromosomes that fail to segregate (E, arrowhead), not observed in wild type (D).

We also analyzed the effect of the *tmi* mutation on microtubule dynamics at the furrow. During furrow maturation in wild type, microtubules of the FMA from opposite sides of the furrow typically exhibit apparent sites of contacts, which also coincide with sites of increased bundling of microtubules at both sides of the furrow (Fig. 4A; Jesuthasan 1998, Wühr et al. 2010, Wühr et al. 2011). Indeed, sites of microtubule bundling at both sides of the furrow at early stages of furrow formation precisely align along the length of the furrow (Fig. 4A’,E), consistent with microtubules from both sides of the furrows interacting at specific sites of bundling along the furrow. In contrast, in *tmi* mutants at the same developmental stage, FMA tubules from both sides of the furrow fail to exhibit the bundling observed in wild type or to contact each other along the furrow (Fig. 4B’,F). Moreover, FMA tips abutting the furrow are tightly apposed in wild type (width of zone of microtubule exclusion: 1.2 μm, SD=0.296, n=7), whereas these are spaced further apart in *tmi* mutants (width of zone of microtubule exclusion: 7.9 μm, SD=1.615, n=7). At later stages of furrow maturation, when FMA bundles are undergoing their characteristic movement towards distal ends of the furrow (Fig. 4C, C’, 11/11), which continues to be reduced in *tmi* mutants (Fig. 4D, D’, 0/23 with distal enrichment).

**Figure 4.**
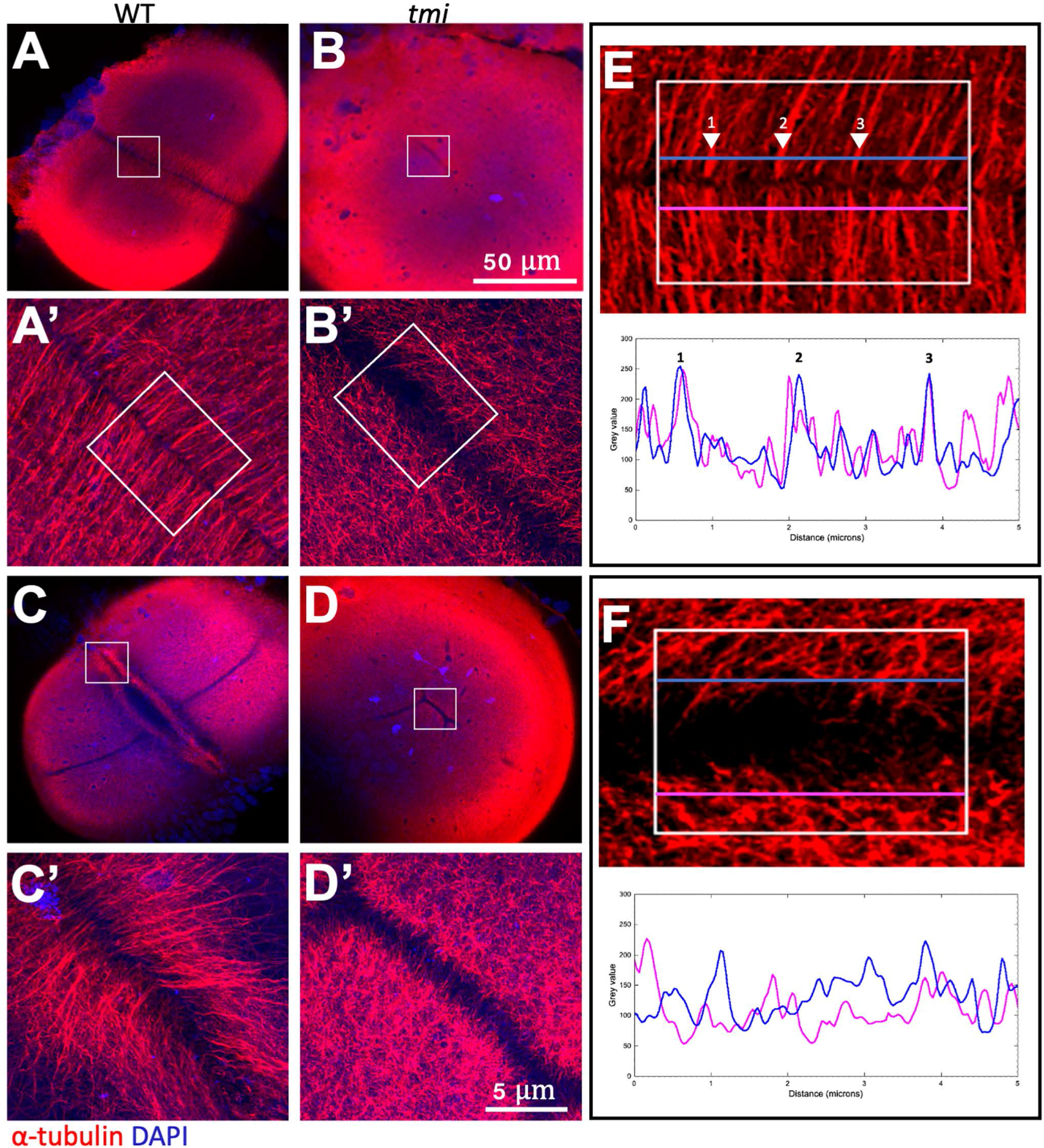
Aberrant clustering and reorganization in the furrow microtubule array. A,B) During early stages of furrow formation, the FMA forms along the length of the furrow, consisting of tubule bundles parallel to each other and perpendicular to the furrow (A, A’). *tmi/prc1l* mutants fail to show a clearly organized FMA and exhibit a reduction in bundling (B,B’, see also Sup. Fig. 6). C-D) Upon furrow maturation, tubules of the FMA reorganize, becoming enriched to the distal ends of the furrow and acquiring a tilted angle pointing distally (C, C’), whereas in *tmi/prc1l* mutants microtubules abutting the furrow remain present along the length of the furrow and do not exhibit tilting (D, D’). E,F) Line scans of FMA bundles along both sides of the furrow show a strong spatial concordance of microtubule bundles in wild type (E, corresponding to insert in A’), consistent with interconnections between bundles across the furrow. Such spatial concordance is absent in *tmi* mutants (F, corresponding to insert in B’). Images are representative of 9 regions of interest for each condition. Numbers on major peaks in (E) correspond to major sites of bundling.

Thus, maternal *tmi/prc1l* function is essential for spindle formation as well as for the bundling and interconnection of microtubules of the FMA along the furrow.

### Requirement for *tmi/prc1l* function during meiosis

Previous studies in zebrafish and other systems have shown that factors involved in early embryonic mitotic cytokinesis also cause cell division defects during the asymmetric cytokinesis that occurs during meiosis (Courtois et al. 2012). We therefore tested for potential meiosis defects in oocytes of *tmi* mutant females through labeling of a-tubulin and DNA. In eggs from control wild-type females 5 minutes post-water activation (mpa), in the absence of sperm, an elongated meiotic spindle with distinct sister chromatids sets can be observed (Fig. 5A, 5/5). In these wild-type water-activated eggs at 8 mpa, this structure transitions into an incipient midbody flanked by DNA that has undergone segregation to the spindle poles (Fig. 5B, 9/9). By 20 mpa, each DNA mass exhibits characteristic behaviors, with the female pronucleus acquiring a decondensed appearance and the polar body a highly condensed morphology, with both masses remaining at this stage attached to the meiotic midbody (Fig. 5C, 7/7). In water-activated eggs from homozygous *tmi* females, a meiotic spindle is visible, although this spindle has a short and tubby appearance (Fig. 5D, 3/3). By 8 mpa in *tmi* mutants, the spindle does not resolve into a midbody-like structure, with sister chromatids unable to undergo segregation to the spindle poles (Fig. 5E, 4/4). Eventually, by 20 mpa both chromatid sets appear to collapse into a single mass of DNA (Fig. 5F, 3/3). This DNA mass, formed after failed meiotic cytokinesis, is predicted to have a diploid chromosomal component and presumably participates in post-fertilization events.

**Figure 5.**
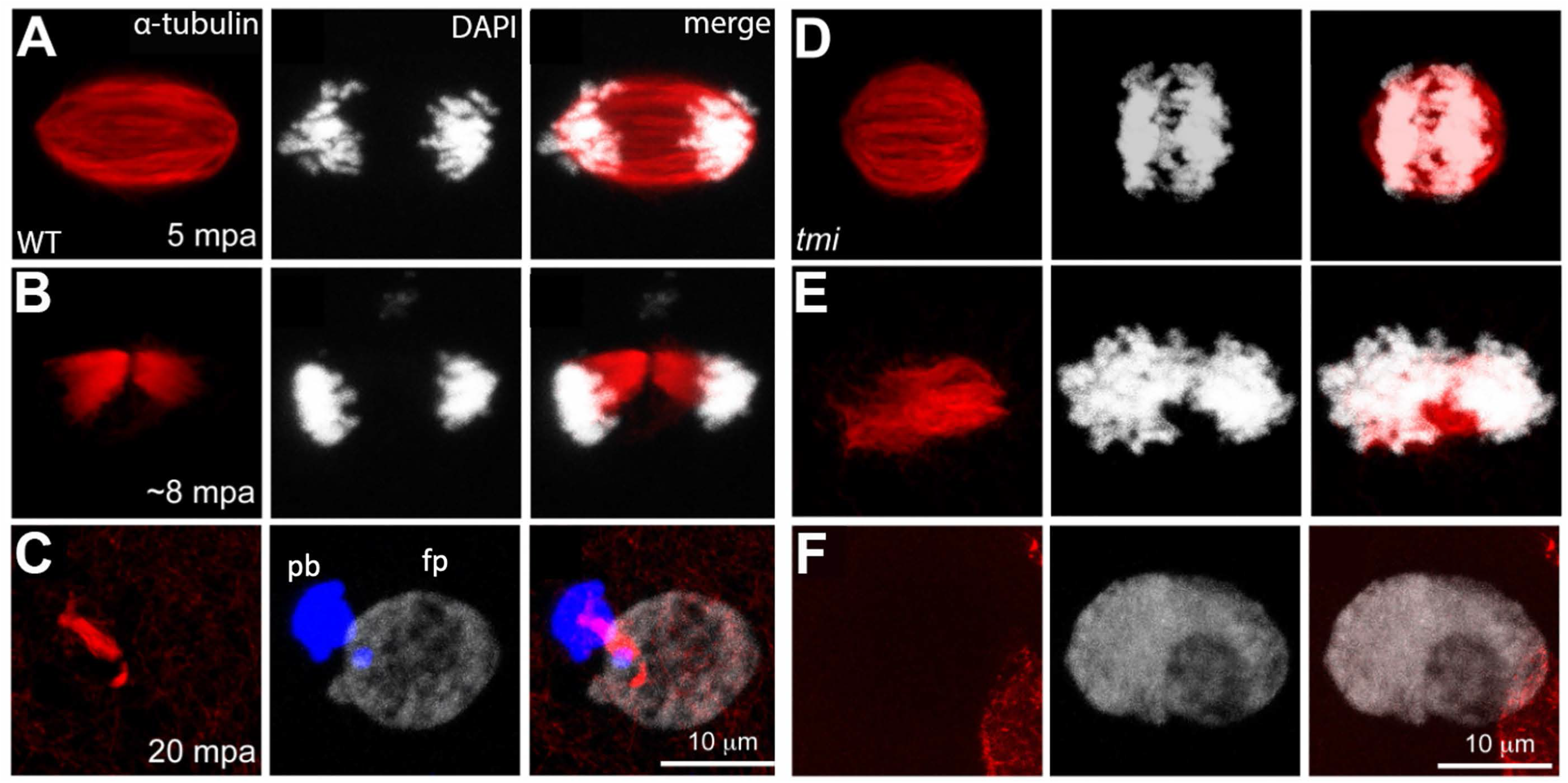
Meiotic spindle defects in *tmi/prc1l* mutants. (A-C) In wild type, spindles for second meiotic division exhibit an elongated shape (A) that resolves into mid-body like structures (B) and becomes highly bundled during abscission (C). During the last stages of cytokinesis, the midbody becomes asymmetric with a longer region in adjacent to the polar body (pb, left DNA mass in C) and a shorter region adjacent to the female pronucleus (fp, right DNA mass in C). Note polar body has become condensed whereas the female pronucleus has undergone decondensation. (D-F) In eggs from *tmi/prc1l* mutant females, spindles appear shorter with less defined poles (D) and fail to form a well-defined midbody (E), leading to failed cytokinesis and a gamete that lacks a polar body and contains a single DNA mass (F), presumably of diploid content. In all panels DNA is represented in white, except the polar body in (C), which has been color coded in blue in the appropriate Z-stack sections to better highlight this structure.

Thus, similar to the embryonic divisions, *tmi* mutants exhibit defects in spindle and midbody organization during meiosis.

### Prc1l is required for vegetal microtubule reorganization involved in axis induction

While carrying out genetics crosses to propagate lines carrying the *tmi* mutation, we noticed that a significant proportion of embryos from mothers heterozygous for the *tmi^p4anua^* mutant allele (genotypically *tmi^p4anua^*/+), display a range of ventralized phenotypes (Fig. 6A) similar to those described for maternal-effect mutations causing axis induction defects, such as *ichabod* (Kelly et al. 2000, Bellipanni et al. 2006), *tokkaibi* (Nojima et al. 2010) and *hecate/grip2a* (Ge et al. 2014). This unexpected observation suggested a potential role for Prc1l function in dorsoventral patterning. We reasoned that the observed partially penetrant defects in embryos from mothers heterozygous for a *tmi* mutant allele may be caused by haploinsufficiency for maternal Prc1l product, and that embryos with fully reduced Prc1l function, i.e. embryos from females homozygous for the *tmi* mutant allele (genotypically *tmi^p4anua^/tmi^p4anua^*), might exhibit a fully-penetrant axis induction phenotype. However, the cell division defect in embryos from homozygous *tmi* mutant females precludes the cellularization of embryos and therefore the assessment of axis induction through morphological landmarks. Thus, we assayed instead defects in axis induction in these embryos using molecular landmarks within the axis induction pathway.

**Figure 6.**
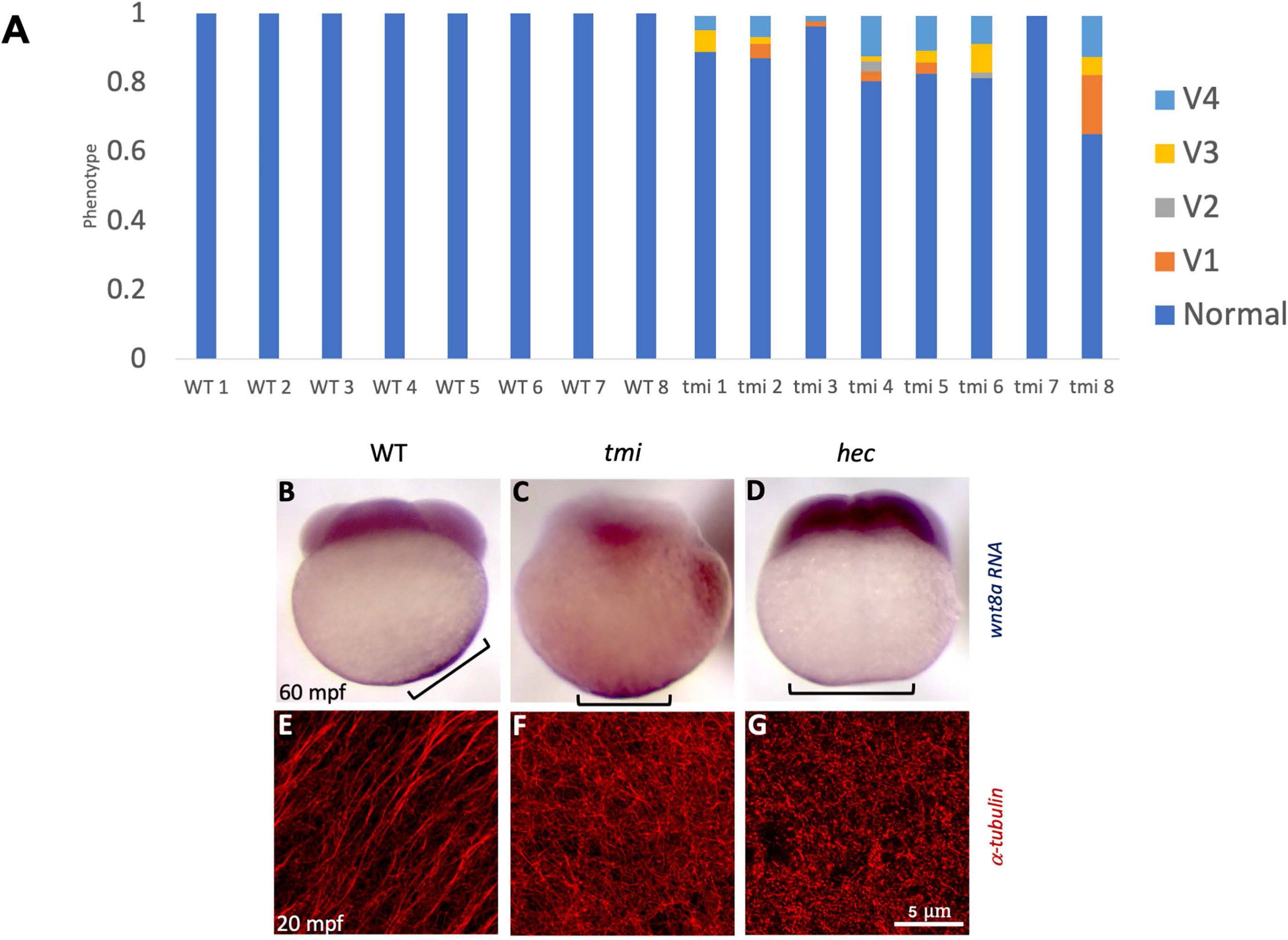
*tmi/prci1l* mutant embryos exhibit axis induction defects. (A) Females heterozygous for the maternal-effect *tmi* mutation consistently produce a fraction of embryos with ventralization phenotypes in 24 hour embryos (V1-V4, ventralization series as in (Kishimoto et al. 1997), with V4 the most severe, radially symmetric phenotype). For each clutch, a minimum of 50 embryos were scored. (B-D) *wnt8a* RNA, originally localized to the vegetal pole of the egg, undergoes a lateral shift towards the future dorsal axis in wild-type embryos (B). This off-center shift is absent in embryos from *tmi/prc1l* homozygous mutant mothers (C), a behavior similar to that caused by the ventralizing maternal-effect mutations *hecate* ((D); Ge et al. 2014)). In (B-D), embryos exhibiting reduced or no label (4-8% of the embryos) were excluded from the scoring. (E-G) Microtubules at the vegetal cortex develop into a parallel array of microtubule bundles aligned toward the future dorsal axis ((E), Jesuthasan and Strähle 1997, Tran et al. 2012, Ge et al. 2014). Vegetal microtubule bundling and alignment is defective in embryos from females homozygous mutant for *tmi/prc1l* (F) and *hecate* (G).

Previous studies have determined that zebrafish *wnt8a* maternal RNA is initially localized to the vegetal pole of the mature oocyte and undergoes a lateral shift after fertilization (Lu et al. 2011), which reflects a symmetry-breaking event that presages axis induction. We used whole mount in situ hybridization to test whether the dorsally-directed lateral shift of *wnt8a* RNA was affected in *tmi* mutant embryos. As expected, wild-type embryos experience an off-center shift of *wnt8a* mRNA toward the presumed dorsal region of the early embryo, observable at 30 and 60 mpf (Fig. 6B, 18/18 with shift). *tmi* mutant embryos at these same stages, however, fail to exhibit an off-center movement of *wnt8a* RNA, which remains instead symmetrically localized at the base of the vegetal pole (Fig. 6C, 0/28 with shift). This RNA redistribution phenotype is similar to that of embryos from *hecate/grip2a* mutant mothers (Fig. 6D, 0/24 with shift; Ge et al 2014).

The off-center shift of *wnt8a* RNA in zebrafish, reminiscent of cortical rotation in amphibians (Houston, 2012), takes place minutes after fertilization and depends on the formation of a parallel array of microtubule bundles at the vegetal pole of the embryo (Jesuthasan and Strähle 1997, Tran et al. 2012, Ge et al. 2014, Welch and Pelegri 2015). As previously reported, vegetal cortex microtubules in wild-type zebrafish embryos begin to undergo bundling and alignment around 14 mpf (not shown), become fully bundled and aligned by 20 mpf (Fig. 6E, 10/10 with microtubule arrays), and start to dissociate at approximately 26 mpf (data not shown; Jesuthasan and Strähle 1997, Tran et al. 2012). In *tmi* mutant embryos at 20 mpf, we find that microtubules at the vegetal cortex do not become aligned into parallel bundles, but instead appear as a disorganized branched meshwork that extends the vegetal cortex (Fig. 6F, 0/12 with arrays, a phenotype again similar to that of *hecate* mutants (Fig. 6G, 0/10 with arrays; Ge et al. 2014). Thus, Prc1l function is essential for the symmetry-breaking event leading to axis induction, likely through a role in the reorganization of vegetal cortex microtubules into aligned parallel bundles.

### Prc1l protein localizes to the spindle, FMA ends and parallel tracks of vegetal cortex microtubules

To determine the subcellular localization of Prc1l we used antibodies against two regions of the predicted 580-amino acid Prc1l protein, corresponding to amino acids 59-102 near the N-terminus and amino acids 242-331 at the mid- to C-terminal region. Western blot analysis showed that antibodies against either protein region recognized a band of the expected size (67 kD) in wild-type embryos that was absent in embryos from females homozygous for the *tmi^p4anua^* mutation (Sup. Fig. 5A and data not shown).

In wild-type embryos immediately after fertilization, Prc1 protein is associated with remnants of the spindle apparatus for meiosis II that begin to undergo bundling (Fig. 7A, 7 mpf, 4/4) and reorganize into compact midbodies (Fig. 7B, 17 mpa, 5/5). Prc1 localization to meiotic midbodies ceases coinciding with the disassembly of the midbody upon completion of polar body extrusion (27 mpa, Fig. 7C, 3/3). In dividing blastomeres of the early embryo, Prc1l protein can be observed to localize both at the metaphase spindle (Fig. 7E, E’, 10/10) and at sites of bundling of the FMA ends along the forming furrow (Fig. 7G, G’, 13/13). Specificity of the antibody labeling is further supported by the absence of label in *tmi* mutants in spindles during meiosis (Fig. 7D, 0/5 with label, compare to A) and embryonic mitosis (Fig. 7H, H’, 0/5 with label; compare to Fig. 7G,G’).

**Figure 7.**
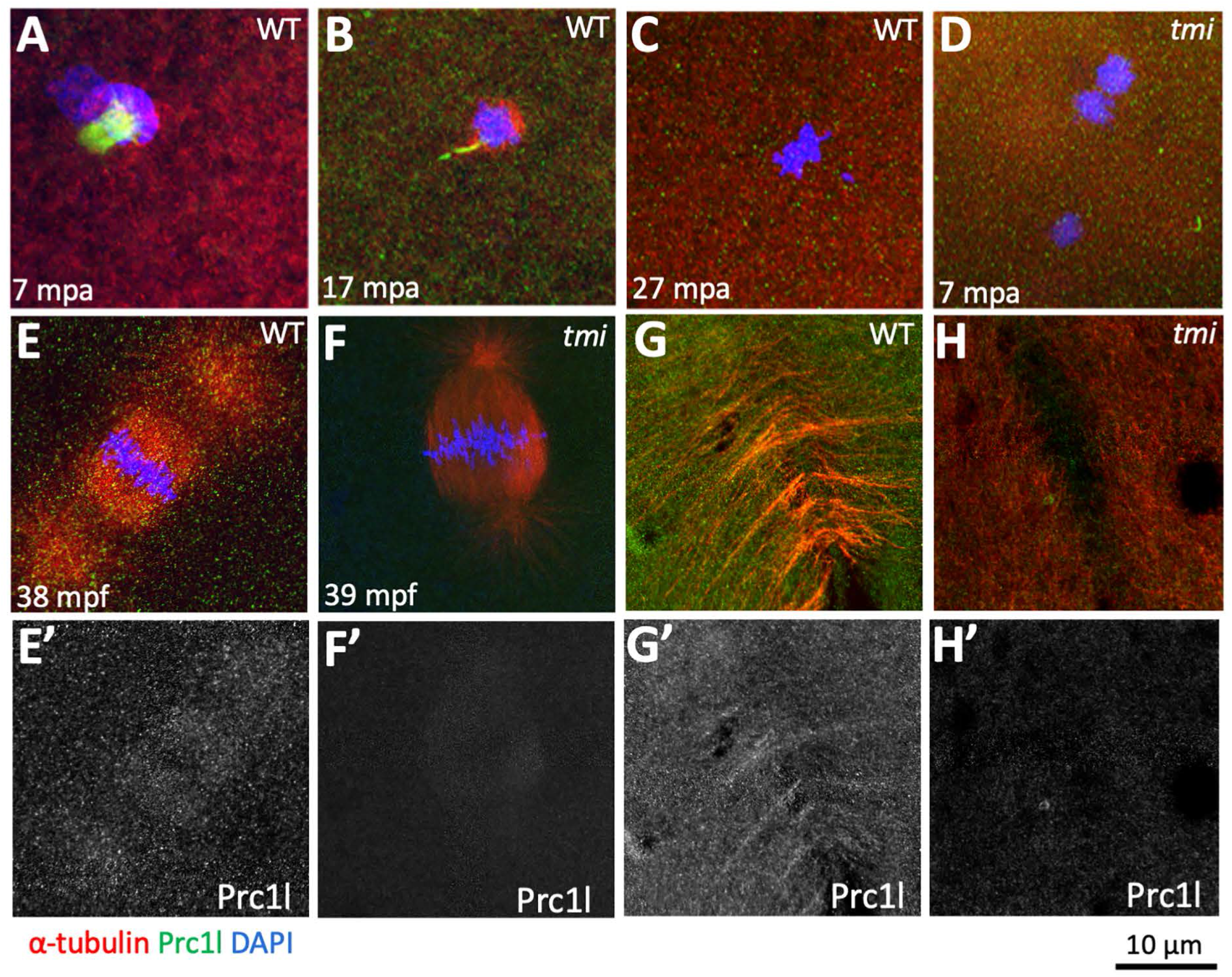
Localization of Prc1l protein to microtubule structures involved in cytokinesis at the animal pole. (A-C) During meiosis II, Prc1l protein localizes to the midbody-like structure associated with the resolution of meiosis II, mirroring the increasing bundling of meiotic midbody microtubules (A,B) and becoming delocalized after polar body extrusion (C). *tmi/prc1l* mutants do not show antibody labeling, as expected from antibody specificity (see also (G,H) and Sup. Fig. 5A). (E-H) During embryonic mitotic divisions, Prc1l protein localizes to the microtubule apparatus (E,E’) and the FMA (G,G’), with the expected absence of label in control *tmi/prc1l* mutant embryos (F,F’,H,H’).

Given the unexpected role for Prc1 function in axis induction we also investigated Prc1l protein localization in microtubules at the vegetal cortex during the first cell cycle interphase (15 mpf). Immunolocalization studies within this region revealed that Prc1l protein is localized along parallel tracks of bundled microtubules (Fig. 8A,A’, 17/17). Interestingly, Prc1l protein along these tracks exhibits a pattern of repetitive enrichments, with an accumulation every 14-28 microns and the most common distance between enrichments at about 19 microns (average = 18.9 μm, SD=4.19, Fig. 8C; Sup. Fig. 5B,B’). Colocalization control images, where the color labeling channels have been rotated by 90 degrees, show no repetitive pattern of Prc1l protein accumulation (Fig. 8D, Sup. Fig. 5C,C’). Both microtubule bundling and Prc1l protein repetitive pattern of accumulation are absent at the vegetal cortex of *tmi* mutants (Fig. 8B,B’,E, 8/8).

**Figure 8.**
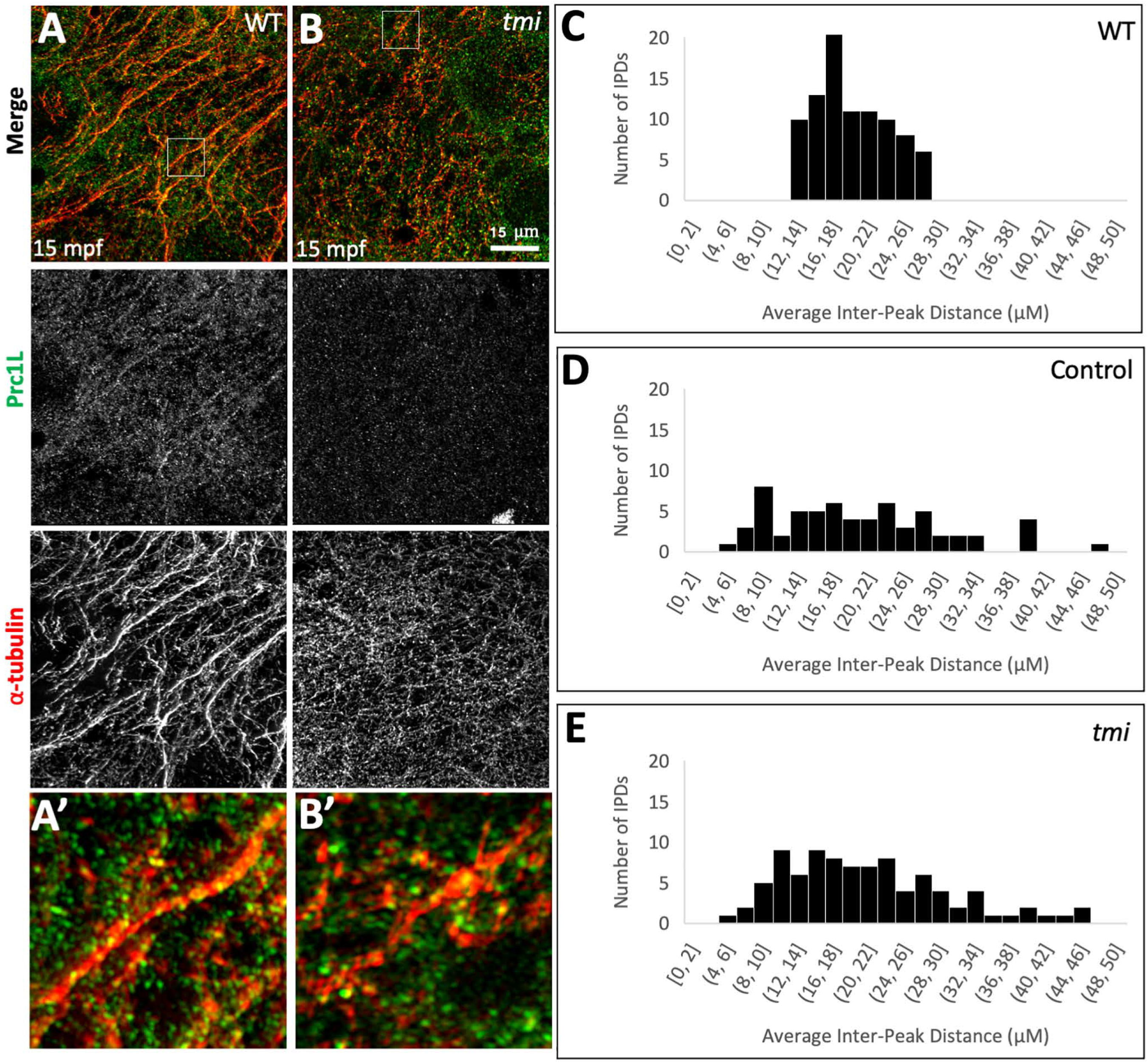
Localization of Prc1l protein to aligned vegetal microtubule arrays at the vegetal pole. (A,B) Array of aligned bundles of vegetal cortical microtubules and localization of Prc1 protein along bundles in wild type ((A), magnification in (A’)) and *tmi/prc1l* mutants ((B), magnification in (B’)). In mutants, microtubules show reduced microtubule bundling and alignment as well as reduced Prc1l protein colocalization to microtubules. (C-D) Distribution of distances between sites of Prc1l localization along microtubule tracks, showing a restricted distribution range in wild type (C), random distribution in a rotated-channel control in wild type (D) and random distribution in *tmi/prc1l* mutants (E). See Sup. Fig. S5 for details on inter-peak distance analysis.

Previous studies have shown microtubule bundling at the midzone involves crosslinking between antiparallel microtubules (Mollinari et al. 2002, Zhu and Jiang 2005, Bieling et al. 2010, Subramanianan et al. 2010, Kellogg et al. 2016), so that localization of Prc1l protein to tracks of vegetal cortex microtubule tracks, shown to be arranged in a parallel conformation (Houliston and Elinson 1991, Schroeder and Gard 1992, Marrari et al. 2003, Tran et al. 2012, Olson et al. 2015, Wijeratne and Subramanian 2018), was unexpected. The vegetal cortex microtubule bundling defect in *tmi* mutants indicates that, in addition to its ability to mediate antiparallel microtubule intercalation required for midzone and midbody formation, Prc1l functions in the bundling of parallel microtubules. Consistent with different mechanisms for bundling, quantitation shows the average thickness of microtubule bundles in vegetal microtubule tracks is approximately 40% wider than bundled FMA tubule tips at the time in which they are expected to undergo interdigitation (Sup. Fig. 6). In spite of divergence of the Prc1l protein sequence compared to that of zebrafish Prc1a and Prc1b and human PRC1 (Sup. Fig. 7), protein modeling does not reveal obvious differences in the predicted three-dimensional structure for these products (Sup. Fig. 8). Rather, this apparent differential activity may rely on additional regulatory interactions (see Discussion).

Thus, Prc1l protein is present in the egg and early embryo in spindles and in areas of active microtubule bundling during cytokinesis, as well as in aligned vegetal cortex microtubules required for axis induction.

## Discussion

We show that the zebrafish maternal effect mutation *tmi* corresponds to Prc1-like, a gene belonging to the MAP65/Ase1/PRC1 family of microtubule associated proteins. Prc1l is essential for meiosis, mitosis and cytokinesis in the early embryo, and unexpectedly also plays a role in the vegetal microtubule reorganization and bundling necessary for dorsal axis induction. Our results provide further evidence for a period of transition at the egg to embryo that encompasses both meiosis and early embryonic mitoses. Moreover, our findings indicate that a common maternal factor, Prc1l, is performing two different functions at the egg-to-embryo transition, both involving microtubule reorganization but in different regions of the egg/embryo and with different purposes: at the animal pole during cytokinesis of meiotic and early embryonic mitotic cell divisions, and at the vegetal pole prior to cell cycling to generate a symmetry-breaking event essential for axis determination.

### The maternal-effect gene *tmi* corresponds to a *prc1* gene duplicate with an essential function at the egg-to-embryo transition

Meiosis and mitosis of early embryos depend critically on the associated steps of cytokinesis. Furrow formation in wild-type zebrafish embryos shows a characteristic pattern, with an initial membrane indentation at the furrow plane, corresponding to incipient furrow formation, followed by full membrane contraction to the base of the blastodisc at the blastomere-yolk boundary (Kimmel et al. 1995). A subsequent step, thought to be associated with the exocytosis of internal membrane vesicles containing cell adhesion components, results in the formation of two closely apposed membranes, or septum, separating daughter blastomeres (Jesuthasan 1998, Pelegri et al. 1999, Feng et al. 2002). Females homozygous for mutations in *tmi* produce embryos that exhibit shallow membrane indentations corresponding to incipient furrows, but which nevertheless fail to undergo full membrane contraction or adhesive septum formation (Dosch et al. 2004, Nair et al. 2013). We find that *tmi* eggs also exhibit defects in cytokinesis for meiosis, with *tmi* mutant eggs exhibit a lack of polar body extrusion. Thus, *tmi* function is required for the progression of cytokinesis in both meiotic and early mitotic cell divisions.

A positional cloning approach finds that the maternal-effect *tmi* mutation is associated with an early truncation of the Prc1l protein, with gene assignment corroborated through non-complementation to newly-generated CRISPR/Cas9 *pcrll* alleles. Phylogenetic analysis indicates that *prc1l* is a duplicated copy of a canonical *prc1* gene that predates the split between amphibian and fish lineages, with expression analysis suggesting that the gene is exclusively maternally expressed. This is in contrast to two other zebrafish *prc1* genes in zebrafish, *prc1a* and *prc1b*, which exhibit both maternal and zygotic expression. Thus *tmi/prc1l* corresponds to a duplicated *prc1* gene that appears to have acquired a specialized function at the egg-to-embryo transition early in vertebrate evolution, and which has been maintained in amphibians and fish lineages. *tmi/prc1l* function is not limited to the meiotic or early mitotic divisions, but rather encompasses both periods. This is consistent with the need to use previously translated product for events associated with egg activation and the earliest embryonic divisions. Thus, as previously found in the mouse embryo (Courtois et al. 2012), a functional transition at the egg-to-embryo transition does not necessarily coincide with the rapid processes of egg activation and fertilization, but appears to be a longer-lasting period, lasting from meiosis to well within the early embryonic cellular divisions.

### Prc1l acts in microtubule reorganization during meiosis and early embryonic mitosis

In many in vivo and in vitro systems, PRC1 is known to localize and bind microtubules at the spindle midzone, where it mediates antiparallel microtubule overlap, and later to the transient midbody (Mollinari et al. 2002, Zhu and Jiang 2005, Bieling et al. 2010, Subramanianan et al. 2010, Subramanian et al. 2013, Kellogg et al. 2016). In zebrafish we find that, in the absence of *tmi/prc1l* function, eggs undergoing meiosis as well as embryonic blastomeres undergoing mitosis exhibit defects in the microtubule apparatus. During meiosis and mitosis, spindles in *tmi* mutants adopt an abnormal morphology, namely a reduced length during meiosis and a lack of the characteristic spindle symmetry during early embryonic mitosis (depicted for mitosis in Fig. 9, center). These results are consistent with recent studies using Xenopus extracts that show the maternal Prc1l orthologue (named Prc1E in this study), together with the kinesin Kif4A, required to regulate the organization of spindle asters and the extent of interdigitation in the early *Xenopus* embryo (Nguyen et al. 2018).

**Figure 9.**
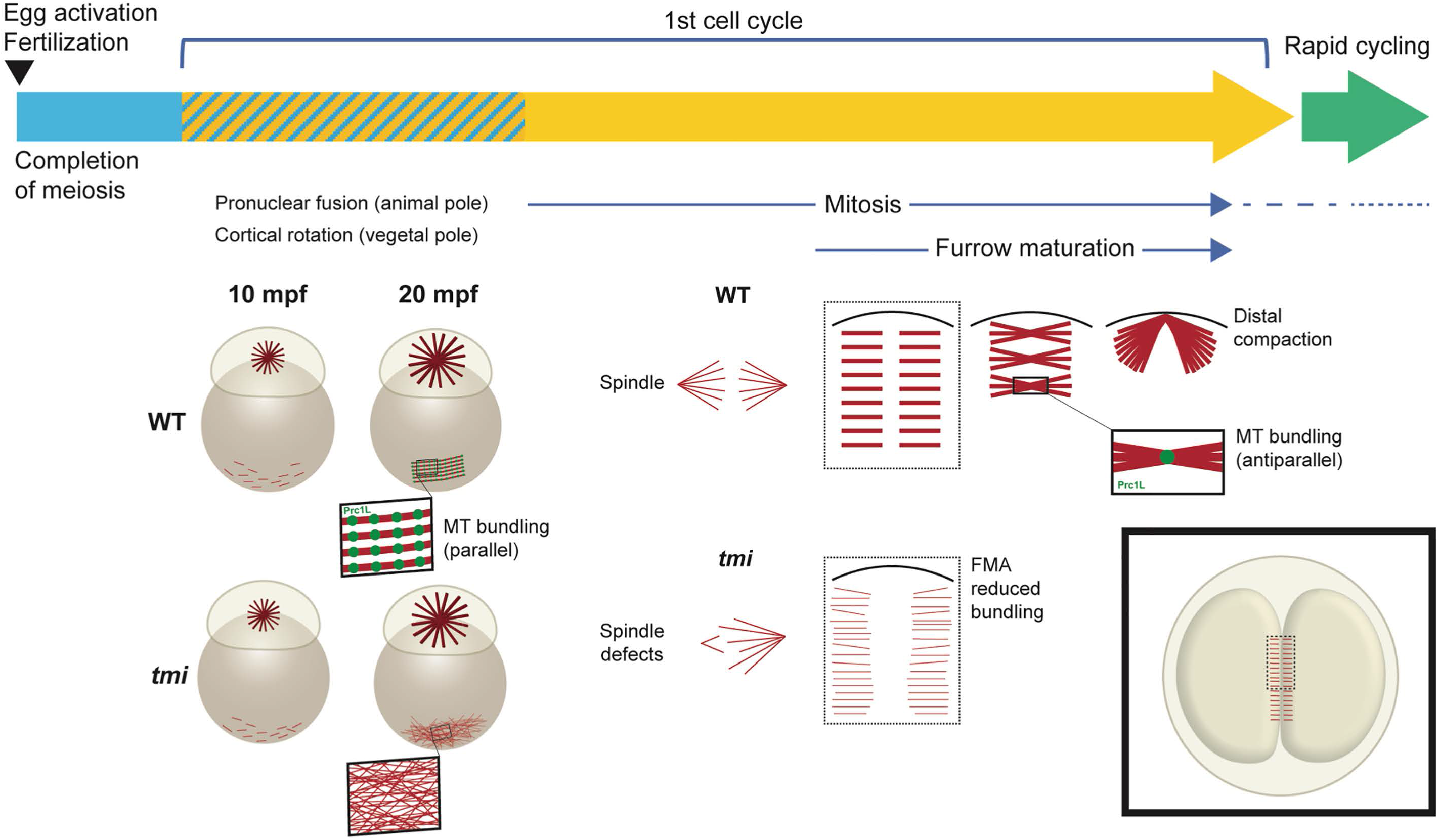
Model diagram highlighting multiple roles for Prc1 at the egg-to-embryo transition. After egg activation and completion of meiosis, the embryo undergoes a first longer cycle, which is followed by rapid mitotic cycles. In wild type, the first cycle includes an interphase period with microtubule growth both at the animal pole, where it forms a sperm aster involved in pronuclear fusion, and at the vegetal pole, where microtubules coalesce into tracks involved in a cortical rotation-like process that generates an early dorsoventral asymmetry. Prc1l is required for the bundling of vegetal cortex microtubules. After the first cycle, the embryo undergoes rapid cell cycling lacking interphase. During this process, Prc1l function is required for microtubule interdigitation at the spindle midzone and the FMA. During cytokinesis, interdigitating microtubules in multiple midbody equivalents gradually coalesce at the distal end of the furrow. Prc1l appears to promote bundling of parallel microtubules during interphase but antiparallel microtubules during mitosis. Not depicted in the diagram, Prc1l is also required for midbody formation during the asymmetric meiotic cell division leading to polar body extrusion.

As the cell cycle proceeds both through meiosis and mitosis, cells undergo cytokinesis through a process that requires the formation of a midbody (Agromayor and Martin-Serrano 2013, Mierzwa and Gerlich 2014). During meiosis, the midbody can be observed to form between the female pronucleus and the forming polar body, a process dependent on factors known to be involved in midbody reorganization such as the Chromosomal Passenger Complex (Nair et al. 2013). Spindles in *tmi* mutant oocytes are unable to form midbodies during meiosis, showing that Prc1l is involved in this process. The large embryonic cells in the early zebrafish embryo, FMA tubules undergo localized bundling at their plus ends along the furrow, and these ends progressively coalesce into a large, compact structure at each of the two distal ends of the furrow (Jesuthasan 1998, Pelegri et al. 1999). These events appear to correspond to a modified process of midbody formation. Indeed, early embryos that lack cues that confer medial-to-distal polarity to the furrow, such as *nebel* mutant or calcium-inhibited embryos, or wild-type blastomeres at later embryonic stages when those cues are no longer present, exhibit bundling of FMA tubules in the medial furrow region in a process indistinguishable from that of a canonical midbody (Eno et al. 2018). In *tmi/prc1l* mutants, FMA formation is defective, failing to undergo microtubule bundling and the subsequent process of distal enrichment. We find that, in wild type when the FMA forms as a parallel array, Prc1l-dependent bundling and crosslinking initially occurs at multiple furrow sites along the furrow, each apparently representing a minimal midbody-like structure. Moreover, these sites of coordinated microtubule bundling appear enriched in Prc1l protein. Previous studies in *Xenopus* have also identified localization of the *Xenopus* Prc1l interacting partner Kif4A at the interdigitating tips of the FMA in this organism, although this study was not able to detect Prc1l labeling at the furrow in embryos, presumably due to an antibody exclusion artefact (Nguyen et al. 2018). Together with previous work (Nguyen et al. 2018), our studies support the notion that Prc1l functions in the cross-linking of antiparallel microtubules across the cleavage plane in the early embryo (Fig. 9, right), similar to the proposed role for this factor at the midzone and midbodies in smaller cell types (Bieling et al. 2010, Agromayor and Martin-Serrano 2013, Mierzwa and Gerlich 2014).

As the FMA undergoes reorganization during furrow maturation, these minimal midbodies migrate and coalesce to form large microtubule masses present at the distal ends of the mature furrow. The directional process that mediates the migration of these bundled ends appear to depend on a wave of intracellular calcium wave traveling along the forming furrow and is dependent on polarized F-actin dynamics (Miranda-Rodríguez et al. 2017, Eno et al. 2018). This alternative mechanism for midbody coalescence at the outward edge of the furrow may reflect an adaptation to cytokinesis completion in large embryonic cells. In zebrafish, ends of the FMA tubules are also associated with germ plasm material (Pelegri et al. 1999, Knaut et al. 2000), a maternal determinant of germ cell fate (Knaut et al. 2000, Hashimoto et al. 2004, Bontems et al. 2009), and this distally-oriented midbody coalescence may additionally reflect the need to promote the tight coalescence of germ cell inducing material for its subsequent internalization by germ cell precursors (Eno and Pelegri 2016).

Thus, defects in microtubule-based structures observed in the blastodisc of *tmi* mutants are consistent with the role for zebrafish Prc1 in the crosslinking of antiparallel microtubule ends, which allows for microtubule bundling and stability of microtubule-based structures during meiosis and the early embryonic mitoses.

### Function of Prc1l in microtubule reorganization involved in axis induction

*tmi* mutant embryos are acellular and lyse at 4-6 hours after fertilization, coincident with the initiation of epiboly in wild type. This acellular phenotype and subsequent lysis would normally preclude observing other potential developmental defects that rely on morphological landmarks for their identification. However, we fortuitously found a role for *tmi* in axis induction by virtue of a partially penetrant phenotype associated with maternal haplo-insufficiency for this gene. Indeed, molecular landmarks show that complete loss of *tmi/prc1l* function results in the absence of the off-center shift characteristic of vegetally localized RNAs and associated with the specification of the embryonic dorsal axis. In wild type, cortical microtubules at the vegetal pole become aligned into parallel arrays, with microtubules becoming visibly thicker as they undergo bundling (Jesuthasan and Strähle 1997, Tran et al. 2012, Ge et al. 2014). In *tmi/prc1l* mutants, this process is affected, with vegetal cortical microtubules exhibiting reduced bundling and remaining largely unorganized, indicating a role for Prc1l in this process (Fig. 9, left). Visualization of the Prc1l protein shows that it localizes to microtubules of the vegetal cortical array, consistent with a role in microtubule bundling. Remarkably, Prc1L protein is found enriched along vegetal microtubule tracks in an apparently repeated pattern, with enrichments at regular intervals of about 19 μm. Previous studies have shown vegetal microtubule tracks as bundles of aligned parallel microtubules, with their plus ends oriented towards the prospective dorsal region (reviewed in Houston 2012). Thus, the localization and function of Prc1L protein is unlikely caused by cross-linking of antiparallel microtubules, as occurs at the spindle midzone and midbody, but is likely instead related to the bundling of parallel microtubules.

Bundling of arrays of microtubules reminiscent to parallel tracks at the vegetal cortex has been previously observed after overexpression of mammalian Prc1 in HeLa cells (Mollinari et al. 2002). In these studies, overexpressed PRC1 could be observed during interphase to both promote and localize to cytoplasmic arrays of long microtubule bundles that are similar to those observed in the vegetal cortex of amphibians and fish. Subsequently, as the cells entered mitosis, overexpressed PRC1 redistributes to the spindle midzone (Mollinari et al. 2002). The cell cycle-dependent effects and localization of overexpressed mammalian Prc1 in mammalian cell lines is therefore remarkably similar to that which occurs for endogenous levels of Prc1l in the early zebrafish embryo. In the latter case, the early embryo undergoes a long first cell cycle with an interphase that includes pronuclear congression and fusion, followed by a period of fast mitotic cell cycles that lack interphase (Kane and Kimmel 1993, see also Tsai et al. 2014). Similarly, early zebrafish Prc1l-dependent cortical microtubule bundles form during the first cell cycle prior to mitotic cycling, and Prc1l-dependent spindle midzone and FMA microtubule interdigitation occurs during the subsequent fast embryonic mitotic cycles. In smaller dividing cells, Prc1 levels are typically low during interphase and rise only during mitosis (Jiang et al. 1998). It may be that high levels of maternally-inherited Prc1 protein present in the egg, and active in the vegetal pole during the first cell cycle interphase, result in the bundling of parallel microtubules, similar to ectopically expressed Prc1 in interphase of smaller dividing cells. The signals that regulate the function of Prc in the bundling of vegetal cortex microtubules remain unknown but may depend on cell cycle-dependent activity of Prc regulators, such as Cdk (Jiang et al. 1998, Mollinari et al. 2002), and/or differential nuclear-cytoplasmic distribution caused by sequestration of factors by the nucleus during the cell cycle. It is also possible that in the early zebrafish embryo regulation of PRC1 activity depends on factors differentially distributed along the animal-vegetal axis. Our studies show for the first time a role for endogenous levels of a Prc1-related product in the bundling of parallel microtubules.

Previous studies suggest a mechanism in which microtubule growth originates local nucleation in multiple locations of the vegetal cortex (Houliston and Elinson 1991, Schroeder and Gard 1992, Marrari et al. 2003, Tran et al. 2012, Olson et al. 2015). Our findings suggest a mechanism in which, following this nucleation, Prc1-mediated microtubule bundling generates and/or amplifies an initial microtubule alignment bias, thus stabilizing microtubule tracks and amplifying a rotation of the vegetal cortex. Loss of function for the maternal Kinesin-1 Kif5Ba results in vegetal cortex microtubule alignment defects (Campbell et al. 2015), and Prc1 has been shown to interact with a subset of Kinesin-family motors (Kurasawa et al. 2004, Zhu and Jiang 2005, Gruneberg et al. 2006, Fu et al. 2009, Bieling et al. 2010, Subramanian et al. 2013, Vitre et al. 2014, Nguyen et al. 2018). Thus, zebrafish maternal Kif5Ba and Prc1l may act in concert to mediate vegetal cortex microtubule alignment.

Additionally, microtubule growth in the vegetal pole in zebrafish (Jesuthasan and Strähle 1997, Tran et al. 2012) occurs at a time concomitant with growth of the sperm-derived monoaster at the animal pole (Dekens et al. 2003, Lindeman and Pelegri 2012), a temporal coincidence that also occurs in Xenopus (Houliston and Elinson 1991, Schroeder and Gard 1992), suggesting embryo-wide signals promote microtubule growth during that early developmental time. Our studies suggest that following such microtubule growth, at vegetal pole growing microtubules are additionally modified through Prc1l-dependent bundling to produce parallel microtubule tracks involved in axis induction. Further research will be required to better understand common signals as well as specific factors involved in these processes.

In summary, we show that a maternal-specific copy of the gene *prc1l* has a role at the egg-to-embryo transition, spanning meiosis and early embryonic mitoses, in key processes involving microtubule reorganization: at the spindle and midbody during cell division, and for vegetal microtubule array reorganization required for axis induction.

## Materials and methods

### Ethics Statement

All animals were handled in strict accordance with good animal practice as defined by the relevant national and/or local animal welfare bodies, and all animal work was approved by the appropriate committee (University of Wisconsin - Madison assurance number A3368-01).

### Fish maintenance and genetic lines

Fish stocks were raised and maintained under standard conditions at 26.5°C. *tmi* was originally isolated from an ENU induced mutagenesis screen. Homozygous *tmi* mutant fish were identified phenotyping, or by using Locked Nucleic Acid (LNA) Endpoint genotyping method that employs 2 fluorescent probes; HEX for identification of the wild-type allele, and FAM for identification of the mutant allele. Mutant embryos were obtained by pairing homozygous mutant females with male siblings. Wild-type controls were obtained from the AB line. Embryos were collected and allowed to develop in E3 embryonic medium until the desired stage, at which point they were fixed using 4% paraformaldehyde. All maternal-effect phenotypes are 100% penetrant with respect to early embryonic lethality (no cellularization followed by lysis) and across multiple tested generations, with experiments analyzing at least three different clutches each from wild-type and mutant females. For LNA genotyping, fish were anesthetized with 0.014% tricaine and the tail fin was clipped using a razor blade and placed in 100μl of 50mM NaOH. Tissue lysates were incubated at 95°C for 20 minutes then cooled to 4°C for approximately 5 minutes. 1/10^th^ the volume of Tris HCl pH 8.0 was added to neutralize the lysate, and it was centrifuged at 13k rpm for 2 minutes. A typical 10μl endpoint genotyping reaction contains 5μl primetime gene expression master mix (IDT), 1μl of 10μM forward and reverse primer mix, 0.5μl, FAM, 0.5μl HEX, DNA between 10 −25ng of genomic DNA and ddH_2_O up to 10μl.

The primers for LNA genotyping of *tmi* were: Primers (5’ -> 3’):

Forward: CTGCTCACGCATGACAATATC; Reverse: AGTTGATAGAGAGTAGAGACACTTT, and the LNA Probes were:

HEX: /5HEX/AGC+C+T+T+GAA+A+CT/3IABkFQ/

FAM: /56-FAM/AGC+C+T+A+GAA+A+CT/3IABkFQ/.

### Cloning and sequence analysis

Segregation analysis was carried out between polymorphic markers as previously described (Nair et al. 2013) and found linkage of *tmi* to markers z25665, z65461 and z9233 in linkage group 21, with recombination frequencies of 3/268, 0/132 and 21/52, respectively (varying numbers of total reflect available polymorphic markers within analyzed crosses). The genetically defined region included seven candidate genes with predicted function in cytoskeletal dynamics and/or cell division: sfswap, FYN binding protein, MOZART, mitotic spindle organizing protein 2B, smarcb1b and two novel proteins zgc:193801 and zgc:86764 present in this interval. Full length cDNAs for each of the seven candidate genes were cloned from mature eggs of wild-type and *tmi* homozygous females, sequenced to identify a potential lesion, and only zgc:86764 contained a non-synonymous mutation.

### Complementation test using CRISPR-Cas9

A CRISPR design web tool, CHOPCHOP, was used to identify a guide RNA site, GTGGACATATGGGACAGCATCGG, immediately upstream of the first predicted protein domain of the predicted Prc1l protein, as identified by Ensembl. The guide RNA template was produced using an annealing/fill-in method as previously described (Gagnon et al. 2014). The guide RNA was synthesized using the MEGAshortscriptTM T7 Transcription Kit and purified using an ethanol/ ammonium acetate protocol. The concentration of the guide RNA was measured by Nanodrop, and its integrity was checked by gel electrophoresis. Single-use aliquots of guide RNAs were stored at −80 °C.

To create the CRISPR-Cas9 mutant zebrafish line, we injected a mixture of guide RNA (200pg/nl total final concentration) and Cas9 protein (PNA Bio; 400 pg/nl final concentration) into one-cell stage AB embryos. To confirm Cas9 activity, a subset of the injected embryos was collected at 24 hpf, DNA was extracted from these embryos and a 95 bp fragment across the guide RNA site was amplified via PCR (F: TGAGATCAATCATGCGATGG R: TTTTTACAGTTTGCATTCTCTCCA) and resolved on a 2.5% agarose gel. If Cas9-mediated genetic edits were created in the somatic cells, the population of DNA that contains a variety of INDELS generates a smear, confirming Cas9 activity (Moravec and Pelegri 2019). Once the *prc1l* F0-injected fish reached sexual maturity, offspring carriers of INDELs were identified using fin clip DNA and the above method for INDEL identification, and the induced mutations were sequenced. The genotype of *tmi^p4anua^/prc1l^uw101^* females was confirmed by testing for each allele independently.

### *in situ* hybridization

Whole mount in situ hybridization, labeled by a standard blue visible substrate was carried out as described previously (Thisse and Thisse 2008), using digoxygenin labeled antisense RNA probes against *wnt8a* and *prc1l* RNAs. To generate the *prc1l* antisense probe, the T7 promoter sequence was attached to a 20bp antisense sequence of *prc1l* cDNA via PCR. Subsequently the PCR product was used as a template for in vitro transcription using T7 polymerase and labeling the transcribed RNA with digoxygenin-conjugated ribonucleotides. A sense control probe was also designed by attaching the T7 promoter sequence to the 20bp sense cDNA sequence. These sequences were also used as primers for RT-PCR and qRT-PCR. The *wnt8a* probe (Lekven et al. 2001) was a gift from the Lekven lab, and generated by linearizing with Ap1I.

### Western Blot

Approximately 200-500 embryos were collected from wild-type and *tmi* mutant fish. Embryos were lysed in RIPA buffer using a 22G syringe. Lysates were centrifuges at 13K rpm for 5 minutes at 4°C to collect debris. Protein concentration was determined and 150 or 300 μg was loaded onto precast 4-15% TGX gels (BioRad) and blotted onto PVDF membranes for 1 hour at 100V at 4°C. Membranes were blocked in 5% milk and blotted with 1:200 mouse-anti Prc1l (Abmart) and 1:5000 anti-mouse HRP (ThermoScientific). Membranes were developed using SuperSignal West Pico (ThermoScientific).

### Immunolabeling

Embryos were dechorionated and fixed in paraformaldehyde and glutaraldehyde, and antibody labeling was carried out as previously described (Lindeman and Pelegri 2012). Antibodies against Prc1l were derived from three separate peptides in each of two different regions of Prc1l, corresponding to amino acids 59-102 and 242-331 (Abmart). Both sets of antibodies yielded similar results in preliminary experiments, and the bulk of the analysis was carried out using antibodies against amino acids 59-102. Other primary antibodies used for immunolabeling of fixed embryos are as follows: mouse anti-Prc1l (amino acids 59-102), 1:400 (Abmart), mouse anti-α-Tubulin, 1:2500 (Sigma T5168), rat anti-α-Tubulin, 1:1000 (Abcam ab6161), rabbit anti-β-Tubulin, 1:1000 (Abcam ab15568). Confocal microscopy images were obtained using a Zeiss LSM 510 for fixed images or Zeiss LSM 780 (for high magnification including super-resolution microscopy) and processed with Fiji. For quantification of the interval between Prc1l protein on vegetal cortex microtubules, for each condition (wild-type, *tmi* mutants, and *hec* mutants) 30 regions of interest (ROI) were randomly placed on the image in Fiji using an adapted version of the macro “RandomSamplePerimeterMethod” (Landini 2008), see also https://blog.bham.ac.uk/intellimic/g-landini-software/. Using the freehand line tool, a line was drawn along the longest microtubule in each ROI. The plot profile tool was then used to assess detection of Prc1l protein aggregate localization to the microtubule. Peaks of colocalized fluorescence were defined as an increase above a grey value of 100. On the plot profiles, the interpeak distances (IPD) were measured between each peak’s apex to determine distance (μM) between protein aggregates. Data presented are averages of IPDs representing the collective IPDs within each ROI. Technical controls were obtained by taking measurements from wild-type images with the green channel (detecting Prc1l protein aggregates) rotated 90° clockwise. To assess relative dispersion of data, the coefficient of variation (CV) was calculated with the “CVTEST” macro on Microsoft Excel using the average IPDs from three images for each condition (90 average IPDs per condition).

### Sequence and protein structure analysis

The DNA and amino acid sequences for the full protein alignment of zebrafish Prc1a, Prc1b, and Prc1l and human Prc1 were collected from Ensembl (http://useast.ensembl.org). Sequence alignment for full proteins and mutations was carried out with PRALINE (CIBVU) and Clustal Omega (Madeira et al. 2019), respectively. The highlighted domains for the Prc1 protein were labeled as previously published (Subramanianan et al. 2010). Gene phylogenetic trees were carried out using Phylogeny.fr (Dereeper et al. 2008, Dereeper et al. 2010).

Three-dimensional protein structures of Prc1a, Prc1b, and Prc1l were predicted using the Phyre2 web server (Kelley et al. 2015). The output secondary structures were color-coated using a rainbow gradient to distinguish the N-terminus and the C-terminus (red to blue, respectively). All predictive protein structures were constructed from a homologous protein folding template, c416yB, provided by the RCSB Protein Data Bank (PDB).

## Supplementary figures

**Figure S1. Conserved protein regions affected in the studied mutations.** A) The *tmi^p4anua^* mutant allele results in the change of a highly conserved leucine (L) at position 221 to a stop codon. B) The CRISPR-Cas9-induced *Prc1l^uw101^* allele contains a deletion of a highly conserved pair of amino acids (GI) near the start of the protein. “*” denotes fully conserved residues, “:” denotes conservation between residues of strongly similar properties, and “.” denotes conservation between residues of weakly similar properties. Amino acids are color-coded according to biochemical properties as follows: red = small, blue = acidic, magenta = basic, green = hydroxyl + sulfhydryl + amine + glycine.

**Figure S2. Non-complementation of the *tmi^p4anua^* and *prc1l^uw101^* alleles.** Embryos from females heterozygous for the *tmi^p4anua^* and *prc1l^uw101^* alleles exhibit fully penetrant maternal-effect phenotypes indistinguishable to embryos from *tmi^p4anua^/tmi^p4anua^* homozygous females. A) Live phenotype shows presence of cells in the 512-cell wild-type embryo but absence of cellular membranes in embryos from transheterozygous *tmi^p4anua^/prc1l^uw101^* females. B) Quantitation of the live phenotype. Data is pooled from embryos derived from 3 wild-type females and 2 *tmi^p4anua^/prc1l^uw101^* transheterozygous females, confirmed through genotyping for both alleles. C) Labeling to detect cell adhesion junction component ß-catenin and DNA (DAPI, blue) shows discrete cell membranes with a nucleus in all cells (some out of focus in this cross-section) in wild type (n=5). In embryos from transheterozygous *tmi^p4anua^/prc1l^uw101^*, there is no ß-catenin accumulation (n=7), reflecting lack of membrane formation, and DNA clusters appear as irregular, larger aggregates.

**Figure S3. Expression pattern of *prc1* family genes.** A) Expression profiles according to available RNA-seq data in the Expression Atlas. *prc1a* is expressed at high levels through mid-somitogenesis, when its levels become reduced. Note *prc1a* expression is present prior to the activation of the genome at the end of the blastula period (3-4 hpf) and persists significantly past that transition, suggesting the RNA for this gene is both maternally inherited and zygotically transcribed. *Prc1b* is found at low or background levels throughout early development. *Prc1l* exhibits an expression profile consistent with a maternally inherited RNA that becomes degraded towards the end of the blastula period. Apparent increases from the zygote to the 2-cell stage are likely artefacts of increased RNA recovery due to RNA polyadenylation after egg activation (see for example Lee et al. (2013)). B) In situ hybridization using antisense probe to detect *prc1l* RNA in wild-type and *tmi* mutant embryos at 45 mpf, showing ubiquitous distribution of maternally inherited Prc1l RNA in wild type and unaffected levels in *tmi* mutants. Sense probe against wild-type embryos was used as a specificity control.

**Figure S4. Gene phylogenetic tree for *prc1* family homologs in selected species.***Danio rerio prc1l* clusters with gene duplicates found in other teleost fish such as *Astyanax mexicanus* (Mexican tetra or blind cave fish), *Salmo salar* (Atlantic Salmon) and *Clupea harengus* (Atlantic herring), but also clusters closely with gene duplicates in anuran amphibians such as *Xenopus laevis* and *X. tropicalis. prc1a* and *prc1b* appear to be recent duplicates within the lineage containing *Danio rerio*, but not present in the other analyzed species, including other teleost fish. The prc1a and prc1b copies in zebrafish are most closely similar to the prc1 copy found across vertebrates including anuran amphibians, the chick, mouse and humans. Xenopus *prc1l* is the same as *prc1E* in Nguyen et al. (2018).

**Figure S5. Analysis of Prc1l protein distribution at vegetal pole microtubules.** A) Validation of anti-Prc1l antibodies using Western blot analysis. A band of the expected molecular weight (ca 67 kD) is found at high levels in extracts of wild-type 1-cell embryos, but is absent in extracts from similarly staged *tmi* mutant embryos. Control detection of actin using an anti-actin antibody shows similar levels of loaded protein. B,C) Immunofluorescence of representative images at the vegetal pole of microtubule tracts (red) and Prc1 protein (green) in wild type, showing the unaltered image (B) and a technical control for colocalization where the green channel is rotated 90° clockwise (C), in this example using the same image as in (B). In both (B) and (C), the Prc1l protein localization pattern was assessed at 30 randomly placed regions of interest (ROI). (B’, C’) To assess Prc1l distribution along microtubules in each ROI, a line is drawn along the longest microtubule track spanning the ROI and fluorescent co-localization (grey value) was assessed by plot profiles. Prc1l localization patterning was discerned by measuring the inter-peak distance of the co-localization detection exhibited in the plot profile. The numbers on the inserts correspond to each randomly assigned ROI. The plot profile shows that the methods are accurate based on the apparent lack of detected co-localization (grey value) in the microtubule track in the technical controls. This analysis was conducted in wild-type and mutant *tmi* embryos to evaluate vegetal pole localization of Prc1l protein to microtubules summarized in Figure 8.

**Figures S6. Analysis of microtubule thickness along the furrow microtubule array and at the vegetal cortex in wild-type and *tmi* mutant embryos.** A-C) FMA microtubules in wild type (A) and *tmi* mutants (B). Microtubules coalesce in thick bundles along the FMA in wild type (A) but exhibit reduced bundling in *tmi* mutants (B). C) Quantification of the average width of microtubule bundle ends in wild type (red, n=30) and microtubule ends in mutant (green, n=30) embryos along the FMA, showing a reduction in the thickness of FMA ends in mutants. (D-F) Vegetal cortex microtubules in wild type (C) and *tmi* mutants (D). The microtubules at the vegetal cortex of *tmi* mutant embryos show reduced bundling (E) compared to wild type (D). F) Quantification of the average width of microtubule tracks in wild type (red) and unbundled microtubules in mutants (green) at the vegetal cortex (n=30), showing reduced thickness in mutants.

**Figure S7. Comparison of Prc1l to the zebrafish and human Prc1 homologs.** A) Multiple sequence alignment comparing Prc1l to related homologs in both zebrafish and humans. “*” denotes fully conserved residues, “:” denotes conservation between residues of strongly similar properties, and “.” denotes conservation between residues of weakly similar properties. Amino acids are color-coded according to biochemical properties as follows: red = small, blue = acidic, magenta = basic, green = hydroxyl + sulfhydryl + amine + glycine. Blue boxed columns highlight conservation of Threonine residues shown to be targets of the cell cycle regulator Cdk (Jiang et al. 1998, Mollinari et al. 2002). B) Percent amino acid identity for the full protein and individual known domains. Analysis and code as in Clustal Omega (Madeira et al. 2019).

**Figure S8. Predictive 3-D Protein Structures of Prc1a, Prc1b and Prc1l.** Phyre2 web server predicted secondary structures of Prc1a (A), Prc1b (B), and Prc1l (C) by alignment to a homologous protein folding template. Orientation of the predicted structures are in the N-terminus to C-terminus direction and annotated by a rainbow gradient from red to blue, respectively. Percent coverage of query sequence to the homologous folding template is provided in each panel. No obvious differences are observed in the predicted structures.

